# The impact of maternal dietary folic acid or choline deficiencies on cerebral blood flow, cardiac, aortic, and coronary function in young and middle-aged female mouse offspring following ischemic stroke

**DOI:** 10.1101/2022.08.23.505040

**Authors:** Kasey Pull, Robert Folk, Jeemin Kang, Shaley Jackson, Brikena Gusek, Mitra Esfandiarei, Nafisa M. Jadavji

## Abstract

**Background and Purpose:** Adequate maternal dietary levels of one-carbon (1C) metabolites, such as folic acid and choline, play an important role in the closure of the neural tube *in utero*; however, the impact of deficiencies in 1C on offspring neurological function after birth remain undefined. Stroke is one of the leading causes of death and disability globally. The aim of our study was to determine the impact of maternal 1C nutritional deficiencies on cerebral and peripheral blood flow after ischemic stroke in adult female offspring.

**Method:** In this study, female mice were placed on either control (CD), folic acid (FADD), or choline (ChDD) deficient diets prior to pregnancy. Female offspring were weaned onto a CD for the duration of the study. Ischemic stroke was induced in offspring and after six weeks cerebral and peripheral blood flow velocity was measured using ultrasound imaging.

**Results:** Our data showed that 11.5-month-old female offspring from ChDD mothers had reduced blood flow in the posterior cerebral artery compared to controls. In peripheral blood flow velocity measurements, we report an aging effect.

**Conclusions:** These results emphasize the importance of maternal 1C diet in early life neuro-programming on long-term vasculature health.

## Introduction

Maternal nutrition during pregnancy and lactation is recognized as a critical factor determining the health of offspring ^1–5^. In addition, early life nutritional cofactors are also critical for fetal development, as well as in determining offspring disease outcome later in life^6, 7^. The Developmental Origins of Health and Disease (DOHaD) theory suggests that prospective chronic diseases are programmed *in utero* ^8–11^. While already compensating for fetal nutrient accumulation and increased maternal metabolic demands ^12^, altered or insufficient maternal nutrition can impact both early development and future offspring health. In offspring, maternal dietary deficiencies have been associated with ventricular septal defects ^13^ and impaired glucose tolerance ^14^, as well as modified neural tube closure ^15, 16^ and neurocognitive development ^17–21^. Beyond this evidence of suboptimal structural development in offspring, maternal nutritional deficiencies have been linked to programming of offspring metabolic ^22^ and epigenetic ^23–28^ adaptations.

Epidemiological studies have demonstrated the effect of maternal diet on lifelong cardiovascular and neurological function ^29–31^. Most of this population-level work reveals an effect of poor maternal health on birthweight and incidence of disease and cardiovascular risk factors in adulthood, such as hypertension and hyperlipidemia ^4, 10^. Such relationships have been shown in numerous global populations and are apparent from birth through early childhood ^32^. Among nutritional cofactors, folates and choline are important players in healthy fetal neurodevelopment due to their involvement in the closure of the neural tube and are components of one-carbon metabolism ^33, 34^. Folates and its chemically synthesized form folic acid are important for fetal neurodevelopment ^35^, as folate requirements during pregnancy are increased by 5-to 10-fold compared to non-pregnant women ^36^. Maternal folates and choline levels during pregnancy have also been shown to be important in the development of the cerebellum and hippocampus ^37^, as well as affecting postnatal myelination trajectories ^38^, short-term memory ^39^, hyperactivity/attention ^40^, neurocognitive development ^41^, and risk of autism spectrum disorder (ASD) ^42^.

Recent work in rodent models has improved mechanistic understanding of how maternal levels of folate and choline impact neurodevelopmental processes ^43^. Akin to human epidemiological studies ^29–31^, murine maternal folate deficiencies have been implicated in adverse reproductive performance, implantation, and fetal growth ^44^. During pregnancy and lactation, maternal dietary folic acid availability has been shown to influence progenitor cell mitosis, and apoptosis in the fetal mouse forebrain ^45^ and hippocampus ^39^. Investigations of maternal perinatal folate deficiencies have revealed reduced hippocampal proliferation, impaired vesicular transport and synaptic plasticity, as well as poor neurite outgrowth ^46^, modified cellular neocortex composition, and diminished complexity and arborization of projection neurons ^47^ in offspring. In addition to these structural observations, both genetic and epigenetic modifications have been observed. For example maternal folate deficiencies has been shown to reduce expressions levels of brain derived neurotropic factor (BDNF) and H3K9me2 (an epigenetic modification to the DNA packaging protein Histone H3) in the fetal hippocampus, while folic acid deficiency for two generations, being reported to significantly enhance *de novo* mutations accumulation during meiosis ^48^.

Choline, another one-carbon metabolite implicated in a number of diverse biological processes ^49^, has yielded variable results in animal models of neurodevelopment. Effects of maternal choline deficiencies, such as defective layering of the cortex, reduced cortical size and brain weight ^50^, and modified hippocampal electrophysiology ^51^ and neurogenesis ^52^, have been observed in offspring. Beyond these physiological and histological findings, choline and folate deficiencies have been shown to elicit similar adverse effects, such as impaired homocysteine re-methylation, oxidative stress, and endothelial dysfunction in murine cerebrovasculature ^53, 54^; effectively demonstrating the link between maternal one-carbon metabolites and typical fetal neurodevelopment.

The link between maternal nutrition and fetal development is abundantly clear, but the long-term (postnatal) effects of maternal nutritional deficiencies on adult offspring are not clearly understood. In this study, we aim to improve our understanding of one-carbon metabolite dietary requirements during pregnancy and its impact on early life programming of adult offspring neurovascular diseases, such as ischemic stroke. Stroke is among the leading causes of death globally and its prevalence as a major health concern is predicted to increase, as the global population ages. Additionally, the demographics of populations change, for example there is a rise in the number of people affected by hypertension, diabetes, and obesity, all increasing risk factors for ischemic stroke^55–57^. Furthermore, in stroke survivors there is an increase in disability, which leads to a reduction in the quality of life for a patient ^58–60^. One of the many reasons these problems exist is that the majority of preclinical studies are targeted only towards male subjects ^61^. Over 90% _o_f preclinical studies use strictly male mice whereas all clinical studies use equal part male and female participants ^61,62^(p201). This makes clinical pharmaceutical findings favor better outcomes in males ^63, 64^. To address the gap in the literature, we investigated the impact of perinatal maternal nutritional deficiencies in folic acid or choline on cerebral blood flow velocity after ischemic stroke in young adult and middle-age female offspring. Since cerebral vasculature blood flow is also directly affected by changes in cardiac and peripheral vascular function and structure ^65–67^, we have further investigated the biophysical properties of the heart and aorta in our experimental groups.

## Methods

### Experimental Design

All methods were performed in accordance with the relevant guidelines and regulations. All animal experimentation was performed following approval by the Midwestern University Institutional Animal Care and Use Committee (Protocol 2983) in accordance with animal welfare and ARRIVE guidelines.

Experimental conditions are summarized in Figure 2. Briefly, for the maternal cohort (n = 30) female and (n = 30) male C57BL/6J mice (RRID: IMSR_JAX:000664, Jackson Laboratories) were purchased from Jackson Laboratories. The mice were acclimatized for one week to controlled housing conditions (22 ± 1°C, 12h-light/12h-dark cycle) with *ad libitum* access to food and water. At two months of age (Day 0), females were randomized to control (CD, TD.190790), and commercially (Envigo) prepared folic acid (FADD, TD.01546) or choline (ChDD, TD.06119) deficient diets and maintained on these diets for four weeks prior to mating, and later throughout pregnancy and lactation (Figure 1). Levels of folic acid, choline bitartrate, and macronutrients in experimental diets are listed in in Table 1 ^39, 53, 54, 68^.

**Figure 1.**
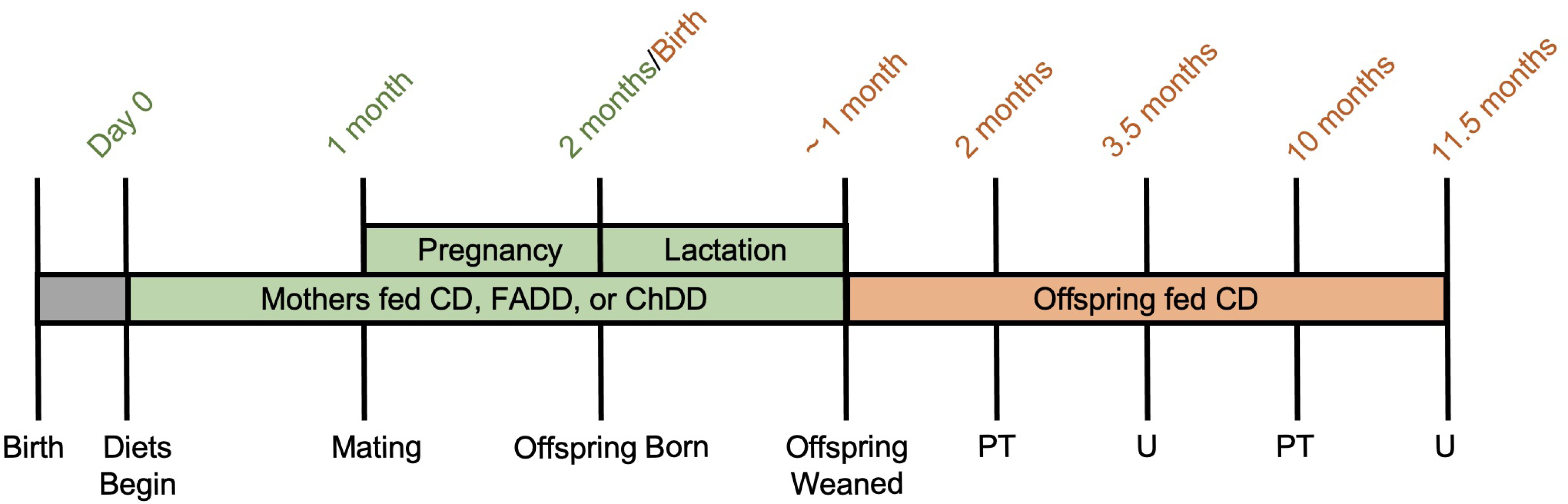
Experimental timeline. Beginning at 2-months-of age female mice were fed either control (CD), folic acid (FADD) or choline (ChDD) deficient diets. The female mice were maintained on these diets throughout the pregnancy and lactation until the offspring were weaned. Once the offspring were weaned, they were fed the CD. Separate cohorts of female offspring at 2 or 10 months of age had ischemic stroke induced via the photothrombosis (PT) model. At 3.5 (CD, n = 6; FADD, n = 7; ChDD, n = 6) and 11.5 (CD, n = 6; FADD, n = 6; ChDD, n = 6) months of age all female mouse offspring underwent ultrasound imaging (U).

**Figure 2.**
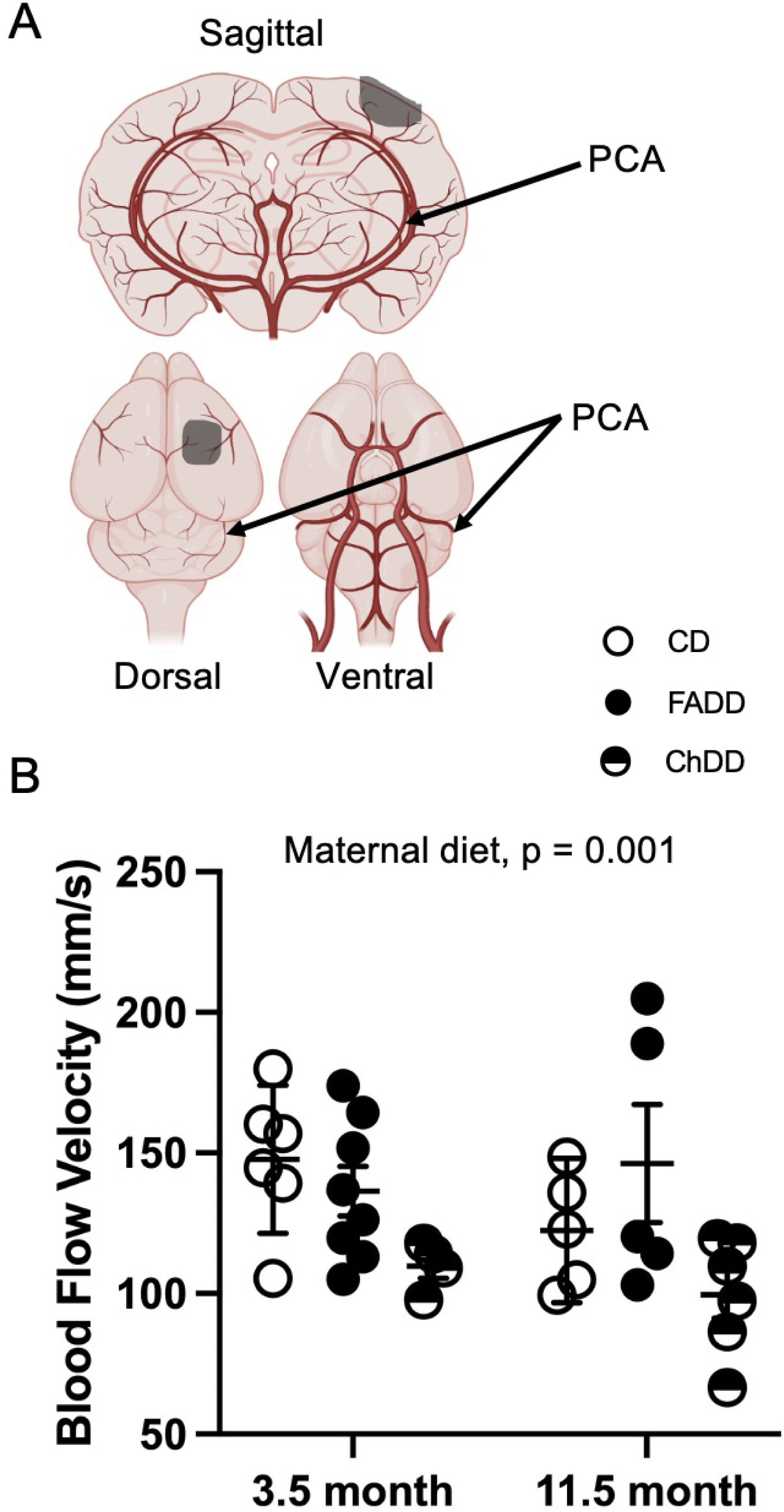
The impact of maternal diet on posterior artery blood flow velocity. **(A)** Visual representation of mouse cerebral vasculature, posterior cerebral artery (PCA) and location of ischemic stroke. **(B)** Blood flow velocity in the PCA after ischemic stroke in 3.5- and 11.5-month-old female offspring from control (CD), folic acid (FADD) and choline (ChDD) deficient diet mothers. Scatter plot with mean ± SEM of 5 to 7 mice per group. * *p* < 0.05, Tukey’s pairwise comparison.

**Table 1.** List of micro- and macro-nutrient contents in Envigo control (TD.01369) folic acid (TD.01546), choline (TD.06119) deficient diets. The Envigo folic acid deficient (TD.95247) was identical to the control diet (TD.04194) except it contained 0.2 mg/kg folic acid. The Envigo choline deficient (TD.06119) was identical to the control diet (TD.01369) except it contained 0.3 mg/kg choline bitartrate.

| Category | Concentration | Category | Concentration | Category | Concentration | Category | Concentration |
| --- | --- | --- | --- | --- | --- | --- | --- |
| **Amino acids** | **(g/kg)** | **Lipids** | **(g/kg)** | **Fat-soluble Vitamins** | **(IU/kg)** | **Nutrient** | **(mg/kg)** |
| Lysine | 14 | Total fat | 60.0 | Vitamin A | 4000 | Choline Bitartrate | 1150 (300) |
| Methionine | 4.7 | Saturated fat | 9.0 | Vitamin D | 1000 | **Fat-soluble Vitamins** | **(IU/kg)** |
| Cystine | 3.6 | Monounsaturated fat | 13.8 | Vitamin E | 75 | Vitamin A | 4000 |
| Arginine | 6.6 | Polyunsaturated fat | 36.6 | Vitamin K | 0.8 | Vitamin D | 1000 |
| Phenylalanine | 8.8 | 4:0 Butyric acid | 0.0 | **Minerals** | **(mg/kg)** | Vitamin E | 75 |
| Tyrosine | 9.2 | 6:0 Caproic Acid | 0.0 | Calcium | 5100 | Vitamin K | 0.8 |
| Histidine | 5.1 | 8:0 Caprylic Acid | 0.0 | Phosphorus | 3000 | **Water-soluble Vitamins** | **(mg/kg)** |
| Isoleucine | 10.1 | 10:0 Capric acid | 0.0 | Potassium | 3600 | Biotin | 0.20 |
| Leucine | 16 | 12:0 Lauric acid | 0.0 | Sodium | 1000 | Choline | 1147.5 |
| Threonine | 7.6 | 14:0 Myristic Acid | 0.0 | Chlorine | 1600 | Folic acid | 2.0 (0.3) |
| Tryptophan | 2.1 | 16:0 Palmitic acid | 6.6 | Magnesium | 515 | Niacin | 30.0 |
| Valine | 11.7 | 16:1 Palmitoleic acid | 0.0 | Copper | 6.0 | Pantothenate | 14.7 |
| Aspartic Acid | 12.1 | 18:0 Stearic acid | 2.4 | Iron | 36.5 | Riboflavin | 6.0 |
| Glutamic acid | 36.6 | 18:1 Oleic acid | 13.8 | Zinc | 35.6 | Thiamin | 4.9 |
| Alanine | 5.2 | 18:2 Linoleic acid | 31.8 | Manganese | 10.5 | Vitamin B<sub>6</sub> | 5.8 |
| Glycine | 3.2 | 18:3 Linolenic acid | 4.8 | Iodine | 0.21 | Vitamin B<sub>12</sub> | 0.03 |
| Proline | 18.1 | Cholesterol | 8.8 mg/kg | Selenium | 0.15 | Vitamin C | 0.0 |
| Serine | 10 |  |  | Molybdenum | 0.15 |  |  |
|  |  |  |  | Chromium | 1.00 |  |  |

At three weeks of age newly born pubs were weaned, and all female offspring were maintained on the CD *ad libitum* for the rest of the experimental timeline. Offspring were randomized to one of two cohorts undergoing photothrombotic (PT) stroke at either 2- or 10-months of age, followed by ultrasound measurements at 1.5 months post-stroke.

### Photothrombosis

When female offspring reached 2 or 10 months of age, ischemia was induced using the photothrombosis model. They were anesthetized with isoflurane (1.5%) in a 70:30 nitrous oxide:oxygen mixture. Core body temperature was monitored with a rectal thermometer (catalog number, 521591, Harvard Apparatus) and maintained at 37 ± 0.2 °C using a heating blanket. 10 mg/kg of the photosensitive Rose Bengal dye (catalog number, 198250, Sigma-Aldrich) was injected intraperitoneally 5 minutes prior to irradiation. A 532 nm green laser (MGM20 (20-25mW, Beta Electronics) was placed 3 cm above the animal and directed to the sensorimotor cortex (mediolateral + 0.24mm) for 15 minutes ^53, 69–71^. At the completion of the study brain tissue was collected, sectioned and stained to confirm ischemic damage ^53, 69–74^.

### Ultrasound imaging

Approximately 1.5 months after ischemic stroke, *in vivo* measurements of posterior cerebral artery (PCA) and aortic function were performed using Vevo® 2100 ultrasound imaging system (FUJIFILM, VisualSonics, Toronto, Canada). Measurements were completed in random order by investigators blinded to treatment groups. The high-frequency, high-resolution ultrasound system used in this study was equipped with a 40 MHz transducer (MS550S) with a focal length of 7.0 mm, frame rate of 557 fps (single zone, 5.08 mm width, B-mode), and a maximum two-dimensional field of view of 14.1×15.0 mm with a spatial resolution of 90 µm lateral by 40 µm axial.

As described in our previous published reports^75–77^, Mice were anesthetized in an induction chamber with 3% isoflurane and 1 L/min flow of 100% oxygen for 1–2 mins, then placed supine on a heated platform and maintained with 1.5–2% isoflurane. Heart rate, electrocardiogram (ECG), and respiratory rate were measured by the four ECG electrodes embedded in the platform. Using a heat lamp and heated platform, body temperature was maintained at 36–38°C and monitored by a rectal probe throughout. The left ventricular (LV) structural and functional parameters, including stroke volume, ejection fraction, fractional shortening, and cardiac output, were calculated from the LV parasternal short-axis M-mode view and recorded at the level of two papillary muscles. An M-mode cursor was positioned perpendicular to the anterior and posterior walls in the middle of the LV for measuring wall thickness. Interventricular septal wall (IVS) thickness during diastole (IVSd) and systole (IVSs) were also obtained from LV parasternal long-axis M-mode view.

Aortic diameters at the annulus, sinuses of Valsalva, and sinotubular junction were measured from the B-mode aortic arch view. Ascending and descending aortic, and posterior cerebral artery (PCA) peak velocities were measured from the pulse wave (PW) Doppler-mode. Aortic pulse wave velocity (PWV) was obtained from the B-mode and Doppler-mode aortic arch view, calculated as PWV (mm/s) = aortic arch distance (d2-d1)/transit time (T1-T2). The PW Doppler mode sample volume was placed in the ascending aorta to verify the time from the onset of the QRS complex to the onset of the ascending aortic Doppler waveform (T1). Using the same image plane, the time from the onset of the QRS complex to the onset of the descending aortic Doppler waveform (T2) was also measured, and the average values for T1 and T2 over 10 cardiac cycles were calculated. Furthermore, the aortic arch distance was measured between the two sample volume positions along the central axis of aortic arch on the B-mode image.

Transcranial Doppler sonography is a non-invasive, non-ionizing, inexpensive, portable, and safe technique that uses a pulsed Doppler transducer for assessment of intracerebral blood flow in the clinical practice ^78, 79^ and has become an important translational tool to evaluate the intracerebral blood flow in animal models. The PCA peak blood flow was measured using the 24MHz (MS250) transducer. The trans occipital window was used to visualize the posterior cerebral arteries and pulsed wave (PW), Doppler-mode was used to measure the PCA peak blood flow velocity.

### Statistics

Ultrasound data was analyzed by two individuals that were blinded to experimental treatment groups using Vevo Lab ultrasound analysis software (FUJIFILM, VisualSonics, Toronto, Canada). Using GraphPad Prism 9.0, two-way ANOVA analysis was performed to assess maternal diet and age effects. Significant main effects of two-way ANOVAs were followed up with Tukey’s HSD post-hoc test to adjust for multiple comparisons. All data are presented as mean ± standard error of the mean (SEM). Statistical tests were performed using a significance level *p <* 0.05.

## Results

### Ischemic damage in brain tissue

To confirm that mice had a ischemic stroke, brain tissue was collected from all animals and damage was confirmed by staining sectioned tissue and quantifying damage volume using microscopy (data not shown) ^80^.

### Cerebral blood flow

After ischemic stroke was induced, approximately 1.5 months, cerebral blood flow was measured in posterior cerebral artery using ultrasound. Representation of posterior cerebral artery is shown in Figure 1A. Maternal diet impacted blood flow velocity within the posterior cerebral artery (Figure 1B; F (_2,28_) = 5.47, p = 0.009). There was significant decrease cerebral blood flow in 11.5-month-old offspring from ChDD mothers (p < 0.05). There was no effect of offspring age (F (_1,28_) = 0.82, p = 0.37.) and no interaction between both factors (F(_2,28_) = 1.25, p = 0.30).

### Measurements of cardiac function and structure

Cardiac function and structure were evaluated at 1.5 months timepoint after ischemic stroke using ultrasound. Both maternal diet (F (_2,28_) = 4.16, p = 0.03) and offspring age (F (_1,28_) = 13.55, p = 0.001) impacted average heart rate (Table 2). Aged (11.5-month-old) offspring from CD (p < 0.01) FADD (p <0 .01) and ChDD (p < 0.01) mothers had higher heart rates compared to 3.5-month-old controls (Table 3). While exclusively offspring age affected ejection fraction (F (_1,29_) = 7.79, p = 0.01) and cardiac output (F(_1,28_) = 35.66, p < 0.0001), there were no pairwise differences (Table 3). There were interactions between maternal diet and offspring age for stroke volume (F(_2,29_) = 4.05, p = 0.03), there was a difference between 3.5- and 11.5-month old FADD offspring (Table 3). There were no differences between groups for fractional shortening and internal diameter in systolic and diastolic measurements.

**Table 2.** Descriptive statistics (Mean ± SEM) and *p*-values results from two-way ANOVA analysis of cerebral and peripheral blood flow in 3.5 and 11.5-month-old female mouse offspring. Maternal diets included a control diet (CD), a folic acid deficient diet (FADD), and a choline deficient diet (ChDD). Mean ± SEM of 5 to 7 mice per group. * *p* < 0.05, ** *p* < 0.01, *** *p* < 0.001 significant diet, age or interaction (diet and age).

| Measurement | Maternal Diet | Offspring Age |  | *p*-values |  |  |
| --- | --- | --- | --- | --- | --- | --- |
|  |  | 3.5 months | 11.5 months | Diet | Age | Diet x Age |
| Average Heart Rate (BPM) | CD | 494.3±13.23 | 524.78 ± 8.82 | 0.0261 * | 0.0010** | 0.1374 |
| | FADD | 504.5 ± 12.68 | 531.42 ± 4.02 | | | |
| | ChDD | 513.0 ± 9.55 | 531.30 ± 6.02 | | | |
| Average Stroke Volume (μL) | CD | 24.04 ± 2.54 | 23.55 ± 4.55 | 0.2744 | 0.2319 | 0.0282* |
| | FADD | 28.67 ± 2.17 | 37.47 ± 5.56 | | | |
| | ChDD | 25.26 ± 2.06 | 38.24 ± 2.08 | | | |
| Average Ejection Fraction (%) | CD | 86.58 ± 4.24 | 84.88 ± 3.55 | 0.1638 | 0.009* | 0.2929 |
| | FADD | 84.68 ± 3.09 | 73.10 ± 4.61 | | | |
| | ChDD | 86.46 ± 2.30 | 74.05 ± 2.49 | | | |
| Average Fractional Shortening (%) | CD | 57.94 ± 6.28 | 53.65 ± 4.18 | 0.3924 | 0.1811 | 0.4998 |
| | FADD | 53.89 ± 3.67 | 53.92 ± 3.88 | | | |
| | ChDD | 54.89 ± 3.13 | 43.70 ± 4.04 | | | |
| Average Cardiac Output (mL/min) | CD | 11.77 ± 1.04 | 20.32 ± 0.78 | 0.7197 | <0.0001*** | 0.5400 |
| | FADD | 14.43 ± 1.08 | 19.91 ± 2.94 | | | |
| | ChDD | 12.94 ± 1.03 | 20.28 ± 0.96 | | | |
| Left Internal Diameter in Systolic (IVSs) (mm) | CD | 0.90 ± 0.08 | 0.90 ± 0.10 | 0.2428 | 0.4138 | 0.3333 |
| | FADD | 1.13 ± 0.10 | 0.91 ± 0.05 | | | |
| | ChDD | 0.95 ± 0.09 | 0.99 ± 0.12 | | | |
| Left Internal Diameter in Diastolic (IVSd)(mm) | CD | 1.37 ± 0.10 | 1.56 ± 0.12 | 0.08098 | 0.5384 | 0.4547 |
| | FADD | 1.71 ± 0.04 | 1.44 ± 0.18 | | | |
| | ChDD | 1.61 ± 0.15 | 1.53 ± 0.12 | | | |

**Table 3.**
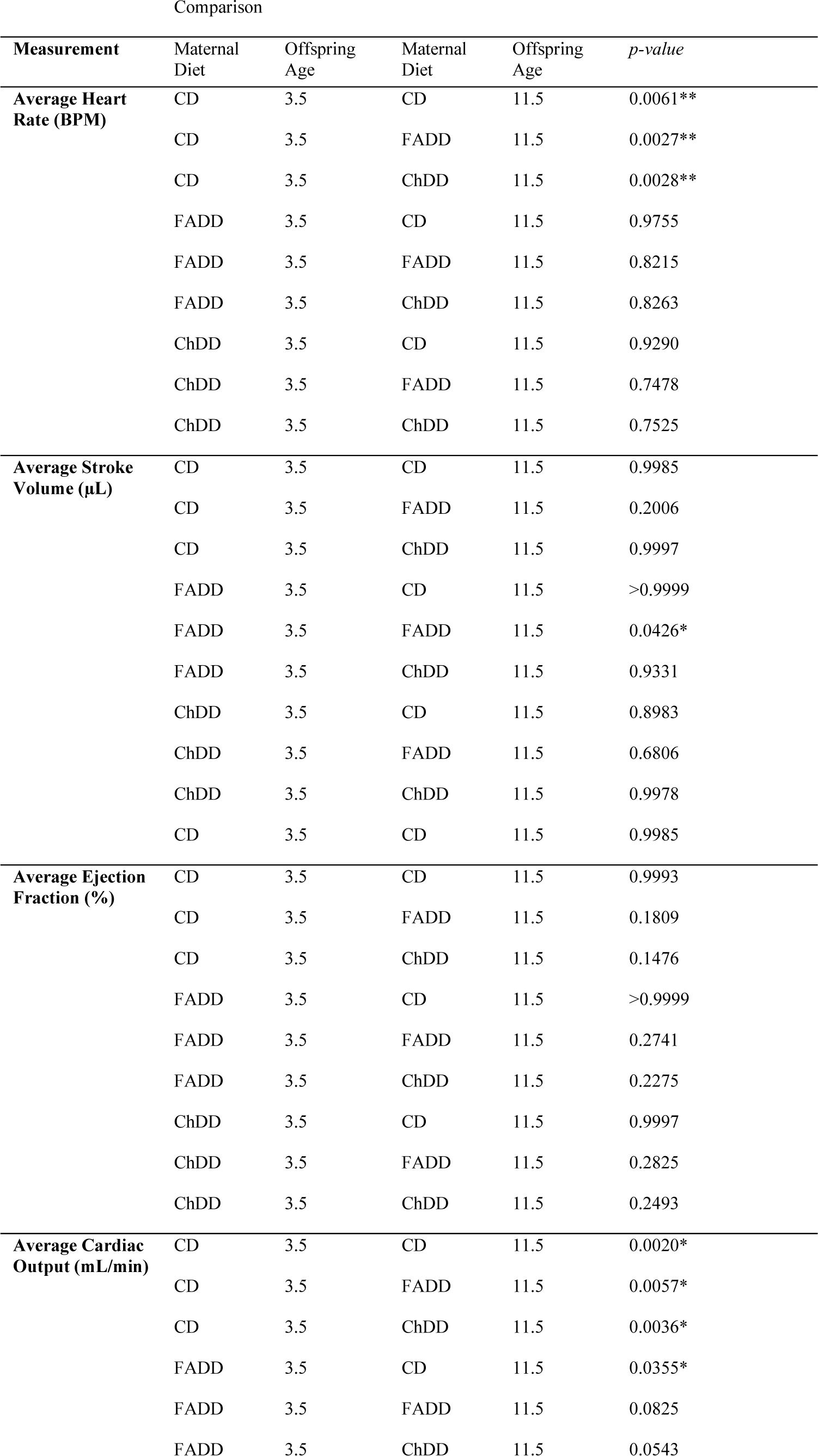
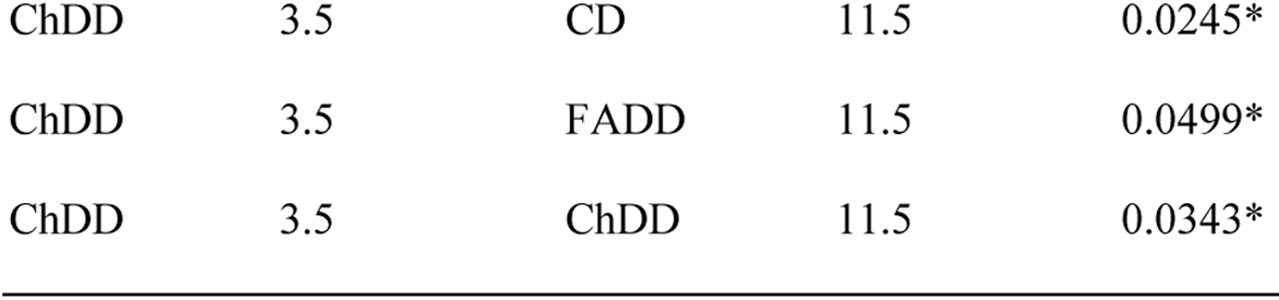
Tukey’s HSD post hoc pairwise analysis of cardiac function in 3.5 and 11.5-month-old female mouse offspring. Maternal diets included a control diet (CD), a folic acid deficient diet (FADD), and a choline deficient diet (ChDD). Presented are the p-values of 5 to 7 mice per group. * *p* < 0.05, ** *p* < 0.01, post-hoc pairwise comparison.

### Aortic pulse wave velocity

Aortic pulse wave velocity is a measure of aortic wall stiffness and was evaluated 1.5 months after ischemic stroke using ultrasound. Offspring age impacted pulse wave velocity (Figure 3A, F (_1,25_) = 33.70, p < 0.0001), there were differences between 3.5-and 11.5-month-old CD (p < 0.01) and ChDD (p < 0.01) offspring. There was no impact of maternal diet (F(_2,25_) = 0.21, p = 0.81) or interaction between maternal diet and offspring age (F(_2,25_) = 2.05, p = 0.15).

**Figure 3.**
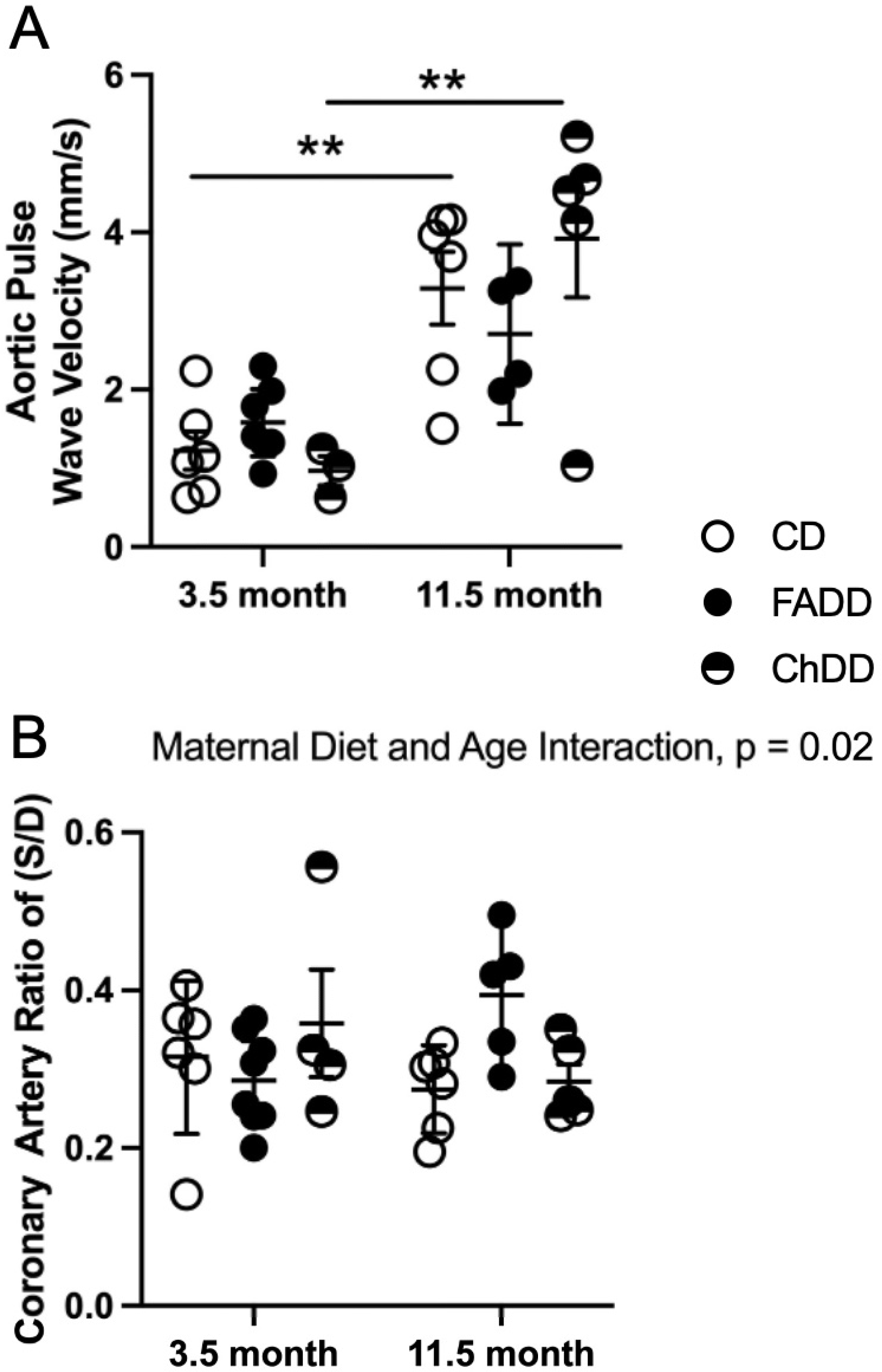
The impact of maternal diet on aorta and coronary artery function. **(A)** Aortic pulse wave velocity as an index of aortic wall stiffness and **(B)** ratio of systolic and diastolic coronary flow velocity after ischemic stroke in 3.5- and 11.5-month-old female offspring from control **(CD)**, folic acid (FADD) and choline (ChDD) deficient diet mothers. Scatter plot with mean ± SEM of 5-7 mice per group. ** p < 0.05, Tukey’s pairwise comparison.

### Measurements of coronary artery function

Coronary artery function was measured using ultrasound 1.5 months after ischemic stroke using ultrasound. Significant interaction between maternal diet and age for coronary artery velocity ratio (Figure 3B, F(_2,28_) = 4.28, p = 0.02). There was no impact of material diet (F(_2,28_) = 0.10, p = 0.38) or age (F(_1,28_) = 0.004, p = 0.95).

## Discussion

The Developmental Origins of Health and Disease (DOHaD) theory suggests that prospective chronic diseases are programmed *in utero*-giving rise to programming of offspring cardiovascular, metabolic, and neuroendocrine dysfunction ^8–11^. Despite impressive evidence of the importance of the maternal environment for fetal growth and development *in utero*, there is not much information available on the impact on offspring as they age, and particularly under neurovascular complications, such as ischemic stroke. Nutrition is a modifiable risk factor for ischemic tissue ^81^ and can also impact outcome after ischemic damage^53, 69–73, 80, 82^. Using an experimental model of ischemic stroke, our study aimed to determine the impact of perinatal maternal dietary deficiencies in folic acid and choline on measures of cerebral, cardiac, aorta, and coronary in offspring, following ischemic injury. Our results demonstrate that maternal diet impact a cerebral blood in offspring after an ischemic stroke in female offspring. The effects of both maternal diet and offspring age, as well as interactions between these variables were observed for numerous cardiac, aortic, and coronary artery function after ischemic stroke in female offspring.

The neurovascular unit (NVU) is comprised of a number of unique neuronal, glial, and endothelial cell types, and recent findings indicate unique cross-talk between neurons and the cerebral vasculature ^83–86^, emphasizing the complex, pivotal role the NVU plays during development and in the progression of neurovascular pathologies like ischemic stroke and cerebral aneurysm ^87–92^. Further, the NVU is responsible for the maintenance of a highly selective blood–brain barrier (BBB) and cerebral homeostasis, as well as the control of cerebral blood flow (CBF) ^93^. The impact of maternal diet on the NVU, modulating integrity of cerebral blood vessels and closure of the neural tube, has been established. Our study adds to these investigations by assessing the response of blood flow within the posterior cerebral artery (PCA) in both young and aged offspring. The contra-lesional PCA was selected as an index of cerebral blood flow due to its spatial and functional independence from the sensorimotor cortex targeted during photothrombotic stroke ^94^, and evident correlation to measures of the Middle Cerebral Artery (MCA) ^95^.

In line with studies detailing the impact of maternal choline on neurovascular development ^16^, our results suggest that maternal choline levels during pregnancy and lactation impair cerebral blood flow in young mice following ischemic stroke. In rodent models, the importance of choline for optimal neurodevelopment is well-established ^96, 97^. Recent work has examined the role of choline in neurovascular interactions as well, modulating levels of anti-angiogenic factors during gestation ^98^, fetal hippocampal angiogenesis ^37^, and promoting the proliferation of rat endothelial cells following hypoxic injury in cerebral vessels ^99^. In this way, choline has been shown to influence neurovascular health across the lifespan and may be implicated in both neurovascular structure and functional response to injury.

The cardiovascular system has also demonstrated effects of choline deficiency, including heart defects ^100, 101^, while higher intake of choline was associated with reduced risk of adult cardiovascular disease ^102^ and amelioration of impaired vagal activity and inflammation in hypertensive rodents ^103^. Our study revealed a diet effect of both maternal choline and folic acid, where deficiencies in either nutrient significantly increased offspring heart rate, regardless of offspring age. While heart rate is a well-known risk factor for cardiovascular disease, our results align well with recent findings associating low heart rate with better functional and cognitive outcomes following ischemic stroke ^104^ and high heart rate with impaired endothelial function and increased ischemic lesion size following stroke ^105^, as well as death due to vascular diseases ^104^. Overall, maternal diet has an established developmental influence on basic measures of cardiovascular and neurovascular health, and may impact offspring programming of the NVU, thereby influencing stroke outcome via endothelial homeostasis via endothelial NO synthase (eNOS) ^106,107^.

We did not observe the effect of maternal diet on cerebral blood flow in 11.5-month-old offspring. This could be due to the well-investigated aging-associated changes in the structural and functional integrity of the vasculature ^65, 108–111^. Therefore, we propose that the difference in effect between young (3.5-month-old) and old (11.5-month-old) offspring is a result of the aging of the control mice. In addition, the presumed damage or endothelial dysfunction induced by the maternal choline and folic acid deficient diets is long-lasting and may contribute to premature aging, generating a mathematically significant difference when compared to young, healthy controls, but only a minor difference when compared to senescent offspring displaying similar levels of vascular dysfunction. This result is supported by the interaction effect of diet and offspring age and requires further investigation. In addition, unique mechanisms drive vascular senescence in males and females ^112^, so our results may be obscured by our study of exclusively female mice. Another interesting result from our study is the significance difference of the coronary artery velocity (S/D) ratio between 3.5 and 11.5-month-old and maternal diet offspring. This result may indicate cardiovascular impairment related to a maternal diet deficient in choline and control diets. However, because the incidence of coronary artery disease (CAD), valvular disease, rhythm disorders, and heart failure increases with age^113^, it appears that folic acid may play a role in programming resistance to this age-related dysfunction.

Outside of heart rate, which is discussed above, age effects were observed for offspring ejection fraction, cardiac output, and pulse wave velocity. Ejection fraction, an index of the left ventricular output, has recently been used as a measure of cardiac mortality risk^114^, with lower percentages indicating cardiac dysfunction. In our study, older mice displayed a significantly reduced ejection fraction, indicating cardiovascular impairment. In a similar manner, our cardiac output data indicate the expected increased cardiac dysfunction as a product of aging^115^. Aortic pulse wave velocity (PWV) was also found to be significantly increased on the older cohort, in line with clinical findings^116^. Controlling cardiovascular risk factors, such as hypertension can decrease risk for ischemic stroke ^117^.

The present study has demonstrated novel data in the field of maternal dietary deficiencies and blood flow after ischemic stroke in offspring. Our data has demonstrated that there are changes in vasculature of female offspring after ischemic stroke, because of maternal diet, this is the first study of its kind. Furthermore, our data points to the need for rodent models spanning a variety of ages for research in age-related diseases such as stroke and vascular dysfunction. We recognize that the exclusion of male subjects in this study may limit our ability to draw conclusions with respect to the impact of sex hormone in observed phenomenon. In future studies, we plan to include male animals, and design experiments that would also allow us to investigate the role of paternal dietary effects. Additionally, we plan to further age animals to ∼20mo after ischemic stroke as well as investigate the role of over supplementation on blood flow after stroke. A detailed analysis of angiogenesis after ischemic stroke might also be prudent.

In conclusion, maternal dietary deficiencies in folic acid and choline have unique roles in cerebral and peripheral blood flow after ischemic stroke in female offspring. Maternal nutrition during pregnancy and lactation has effects, even after infancy and childhood. Our work demonstrated an age effect in animal models encourages further comprehensive longitudinal time-point studies that includes older age animals.

## Acknowledgments

Figure 2A was created with BioRender.com

